# InfoMuNet: Information-theory-based Functional Muscle Network Tracks Sensorimotor Integration Post-stroke

**DOI:** 10.1101/2022.02.10.479324

**Authors:** Rory O’Keeffe, Seyed Yahya Shirazi, Seda Bilaloglu, Shayan Jahed, Ramin Bighamian, Preeti Raghavan, S. Farokh Atashzar

**Affiliations:** Department of Electrical and Computer Engineering, New York University, New York, NY, USA; Department of Medicine, New York University Langone Health, New York, NY, USA; Office of Science and Engineering Laboratories, Center for Devices and Radiological Health, United States Food and Drug Administration, Silver Spring, MD, USA; Departments of Physical Medicine and Rehabilitation and Neurology, Johns Hopkins University School of Medicine, Baltimore, MD, USA; Department of Mechanical and Aerospace Engineering, New York University, New York, NY, USA

## Abstract

Sensory information is critical for motor coordination. However, understanding sensorimotor integration is complicated, especially in individuals with nervous system impairment. This research presents a novel functional biomarker, based on a nonlinear network graph of muscle connectivity, called InfoMuNet, to quantify the role of sensory information in motor performance. Thirty-two individuals with post-stroke hemiparesis performed a grasp-and-lift task while muscle activities were measured using eight surface electromyography (sEMG) sensors. Subjects performed the task with their affected hand before and after exposure to the sensory stimulation elicited by performing the task with the less-affected hand. For the first time, this work shows that InfoMuNet robustly quantifies functional muscle connectivity improvements in the affected hand after exposure of the less-affected side to sensory information. >90% of the subjects conformed with the improvement resulting from this sensory exposure. InfoMuNet also shows high sensitivity to tactile, kinesthetic, and visual input alterations at the subject level, highlighting the potential use in precision rehabilitation interventions.

## Introduction

Stroke is a leading cause of long-term disability worldwide. As many as 76% of individuals with stroke experience upperlimb motor impairment at the stroke onset^1^. This impairment results from disrupted afferent and efferent neural transmission to and from the central nervous system (CNS), ultimately causing delayed initiation and termination of muscle contraction^2^, slowness in developing forces, and disrupted processing and responsiveness to sensory feedback^3^. The collective firing of alpha motor neurons in the spinal cord activates motor units in the muscles, and their recruitment can be examined using surface electromyography (sEMG) to evaluate alterations in motor control^4, 5^. However, there are a variety of muscle activation parameters that can be altered as a result of stroke, which makes it difficult to find a parsimonious and consistently robust method to objectively evaluate changes in muscle coordination and motor control across individuals with stroke.

Stroke-induced motor impairments are more apparent in the contralesional limbs (affected side) than in the ipsilesional limbs (i.e., less-affected side)^6^. In addition, the location of the stroke lesion in the brain can have specific effects on motor and sensory processes^7, 8^. For example, a stroke in the right frontal region may impair temporal motor control on both the affected and less-affected sides, whereas a left parietal stroke may affect the spatial accuracy of upper limb reaching tasks^8, 9^. Specific stroke locations can produce selective visual, tactile, or proprioceptive sensory deficits, which also affect motor planning and performance^10, 11^. However, the less-affected side may provide critical sensory inputs to improve motor planning and performance on the affected side during functional tasks^12^.

Rehabilitation after stroke is personalized based on the type and severity of the deficits and rehabilitation goals of the individual^13^. Quantifying the extent and severity of the impairments using standardized tools such as the Chedoke-McMaster stroke assessment and the Fugl-Meyer Scale can classify patients based on the level of impairment and track improvement with rehabilitation^14, 15^. However, these tools do not quantify sensorimotor integration and are not sensitive to subtle changes in functional motor coordination and control that may occur within a single session. Thus, there is an unmet need for objective functional biomarkers to track subtle changes in motor control, for example, in response to sensory feedback.

The spectrotemporal features of sEMG can, in theory, be the basis for a biomarker to monitor changes in motor control after stroke. This is because stroke alters the fluency of communication between the CNS and muscles, resulting in changes in sEMG activation patterns for everyday tasks^16^. Quantifying changes in sEMG using classical metrics such as the root mean square (RMS) amplitude^16, 17^ and power spectral density (PSD)^5, 18, 19^ of muscle activity can be used to determine the intensity of muscle activation and the use of compensatory strategies. However, these metrics do not offer a means to combine the information from multiple muscles in a task-specific manner to assess coordination.

More advanced methods of sEMG processing such as low-frequency muscle synergies have been proposed to model coherence across muscle groups which play an imperative role in functional task performance^20–22^. However, conventional muscle synergy analysis of the affected and less-affected limbs has shown similar patterns despite clear differences in motor performance. This suggests that muscle synergies may reflect spinal neural control^23^ and may not provide the differential power needed to evaluate the shades of functional impairment caused by alterations in supraspinal neural control.

A more recent method for assessing functional connectivity across muscles is by computing the intermuscular coherence network which is a measure of the degree of information sharing across muscles necessary for sensorimotor control of movement^24, 25^. In this regard, a recent study on postural balance tasks showed that humans reorganize coherence-based muscle networks across limbs in the beta-to-gamma bands (20-60 Hz)^26^. Coherence analysis across various muscle groups may represent the common input from the upper motor neurons to alpha motor neurons^27^. This concept has been tested using invasive needle electromyography as a potential biomarker of upper motor neuron involvement in motor neuron disease^28^. The connectivity between muscles at multiple distinct frequency bands demonstrates how muscle networks can be used to investigate the neural circuitry of motor coordination and provide insights about subtle motor control strategies that cannot be differentiated with other metrics, including muscle synergies^29^.

While the recently-accelerated muscle connectivity research has focused on the use of linear measures of same-frequency coherence (such as delta-band source to delta-band target), cross-frequency connectivity and nonlinear coupling may provide a holistic insight into distributed cortical motor control (motivated by studies on corticomuscular connectivity^30, 31^). Such nonlinear coupling may provide more robust metrics to quantify the effects of stroke and of specific interventions on motor control and coordination. To the best of the authors’ knowledge, the use of a nonlinear measure to quantify a functional muscle network has not been reported in the literature.

Thus this paper focuses on the use of the functional muscle network as a potential biomarker for sensorimotor integration after stroke. We investigate a novel method for assessing temporal connectivity across 16 upper limb muscles (8 in each arm) using a network of nonlinear information-theory-based measure of intermuscular coupling, called InfoMuNet, derived from processing the full-spectrum of sEMG recordings (Fig. 1a,b).

**Figure 1.**
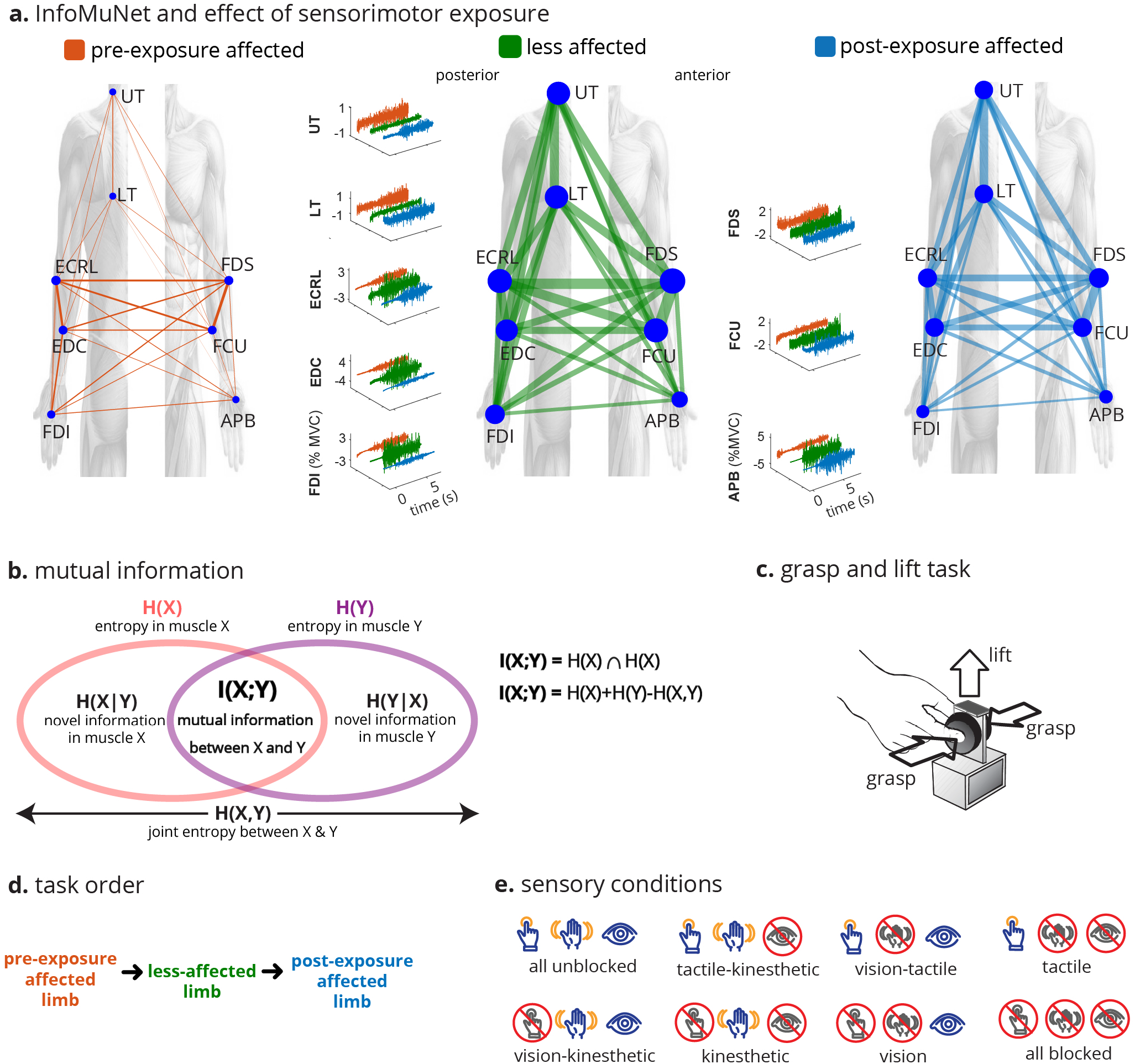
Grasp-and-lift task and InfoMuNet overview. **a.** The median InfoMuNet (i.e., the connectivity network of median mutual information between muscle pairs) across all subjects and trials indicates that the less-affected limb has the strongest connectivity during the grasp-and-lift task. The post-exposure affected shows stronger connectivity trends than the pre-exposure affected limb, highlighting the potential effect of sensorimotor exposure. sEMG plots are representative activations for each muscle. **b.** Mutual information *I*(*X*;*Y*) between two muscle signals is computed using the individual entropies *H*(*X*),*H*(*Y*) and joint entropy *H*(*X*,*Y*). *I*(*X*;*Y*) was quantified between all pairs of muscles signals to form the InfoMuNet on each trial. **c.** For each trial, subjects grasped the instrument with their thumb and index finger and lifted the instrument. **d.** The experiment was completed in two separate sessions. Subjects performed the task with the affected limb only (pre-exposure affected) in the first session. Subjects performed the task first with the less-affected limb followed by the affected limb (post-exposure affected) in the second session. **e.** The task was performed under eight sensory conditions. Subjects completed 14 trials for each sensory condition for a total of 112 trials (14 trials x 8 sensory conditions =112 trials).

Thirty-two individuals with chronic post-stroke upper-limb hemiparesis (21 males and 11 females, age: 57.9 ± 12.7 years, mean time since stroke: 46.4 ± 57.5 months, 15 with right hemiparesis and 17 with left hemiparesis) participated in a multiple-session grasp-and-lift experiment (Fig. 1c) during which their sEMG was recorded from 16 muscles, 8 on each limb. In previous work, we showed that fingertip force coordination in the affected hand changes after exposure to the same task using the less-affected hand^12^. Here, using sEMG, we quantify how sensorimotor exposure to the less-affected limb changes functional muscle coordination in the affected limb. The protocol utilizes alternating hand training (AHT), where the subject first performs the task with the less-affected hand to get exposed to sensory stimuli and then conducts the task with the affected hand (Fig. 1d).

The usefulness of the InfoMuNet approach is compared with that of the classical spectral and temporal metrics obtained using sEMG. The spectral and temporal features of the muscle activity signal (including RMS and PSD of the 16 sEMG channels) are compared with the degree, weighted clustering coefficient (WCC), mean shortest path and global efficiency of InfoMuNet, across the affected and less-affected limbs and various sensory conditions.

We hypothesized that 1) the connectivity metrics derived from InfoMuNet would sensitively capture changes in distributed muscle coordination in the affected limb after contralateral sensory exposure, whereas classical measures (i.e., RMS and PSD of sEMG) would not provide the needed separation power. In other words, although the absolute change in muscle activation may follow a heterogeneous pattern, the information sharing between muscles (that highlights coordinated projections of the CNS) shows an improved pattern after contralateral sensory exposure.

Moreover, we hypothesized that 2) InfoMuNet would show a discriminative sensitivity to contralateral sensory exposure to tactile, kinesthetic, and visual sensory inputs during task performance.

## Results

This study evaluates changes in sensorimotor integration during a functional grasp-and-lift task before and after exposure to the less-affected limb post-stroke. Thirty-two subjects with chronic post-stroke hemiparesis performed a grasp-and-lift task (Fig. 1c) under eight different sensory conditions where visual, kinesthetic and tactile inputs were varied in a random manner (Fig. 1e) over two sessions. At the first session, the task was performed with the affected hand only while sEMG was recorded from eight muscles in the affected limb (preexposure limb). In the second session, subjects performed the task first with the less-affected hand (exposure to preserved sensorimotor information) immediately followed by the affected hand (post-exposure limb).

Subjects performed 112 trials each with the pre-exposure affected limb, post-exposure affected limb, and less-affected limb (14 trials × 8 sensory conditions =112 trials). For each sensory condition, two sets of 7 trials were performed, using a light and a heavy weight. The choice of the light and heavy weights was randomized between 8 different weight combinations ranging from 250-425 (light) and 500-675 g (heavy) in 25 g increments. We used the data from all the weight pairs for all subjects as one group because the preliminary analysis of the sEMG data did not indicate significant differences between light and heavy weights for the metrics used in this study.

Eight bipolar sEMG sensors (Delsys Inc, Natick, MA) were placed on the following muscles on each of the affected and less-affected limbs (Fig. 2a) over the muscle belly of the following muscles: Abductor Pollicis Brevis (APB), First Dorsal Interosseous (FDI), Flexor Digitorum Superficialis (FDS), Extensor Digitorum Communis (EDC), Flexor Carpi Ulnaris (FCU), Extensor Carpi Radialis Longus (ECRL), Upper Trapezius (UT), and Lower Trapezius (LT). Comparisons were made in the muscle activity of the pre-exposure affected, post-exposure affected and less-affected limbs, considering the full task duration.

**Figure 2.**
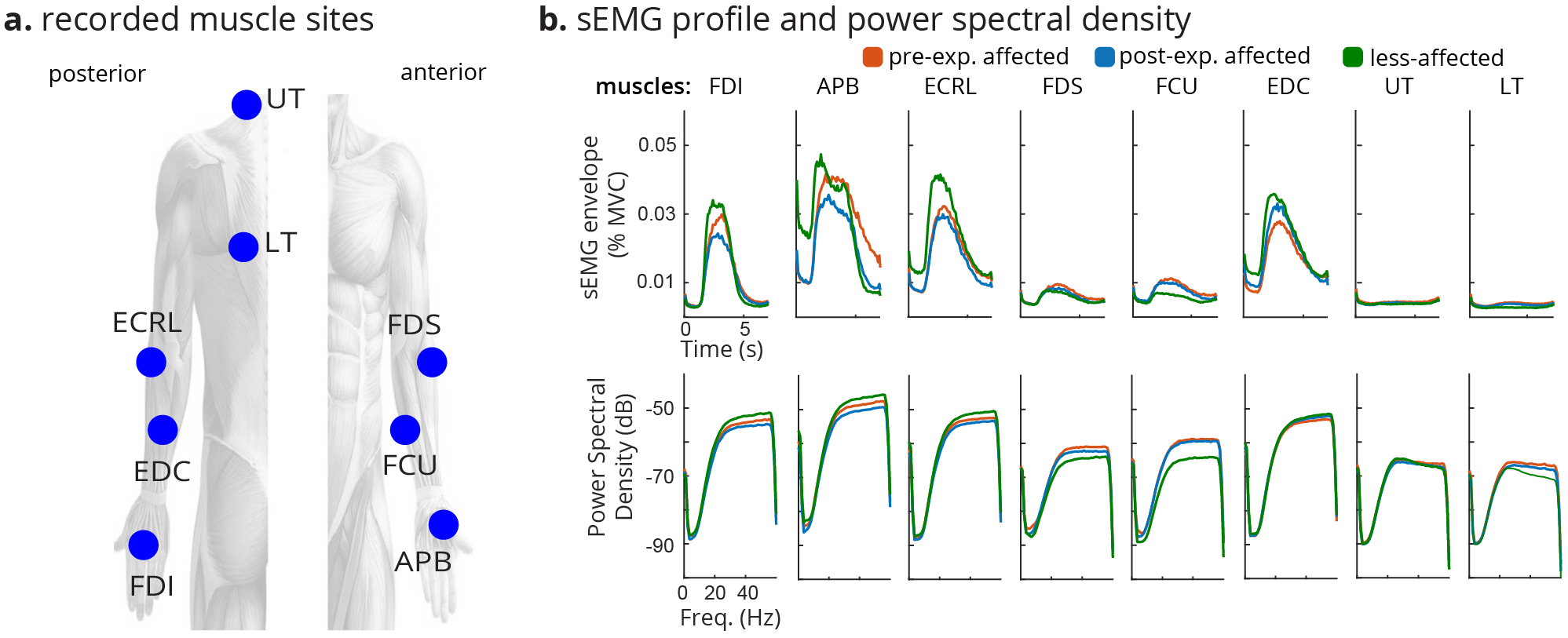
sEMG overview. **a.** Electromyographic signals were recorded from eight muscles on each limb: Abductor Pollicis Brevis (APB), First Dorsal Interosseous (FDI), Flexor Digitorum Superficialis (FDS), Extensor Digitorum Communis (EDC), Flexor Carpi Ulnaris (FCU), Extensor Carpi Radialis Longus (ECRL), Upper Trapezius (UT), and Lower Trapezius (LT). **b.** The median of rectified sEMG envelope (top) across all trials of all subjects for each muscle and the median of PSD of sEMG (bottom) across all trials of all subjects for each muscle are shown. Both the rectified sEMG and PSD varied across muscles and had different trajectories from pre-exposure affected to post-exposure affected, to less-affected limbs.

It should also be highlighted that for a clinically transferable robust sEMG-based biomarker, a linearly-trended behavior would be needed that monotonically changes going from preexposure affected, to post-exposure affected, to less-affected limbs in a consistent direction. This monotonicity is critical if it is claimed that a biomarker can detect subtle changes in the recovery process.

### Classical Spectrotemporal sEMG Analysis

The classical spectral and temporal analyses of muscle activity using rectified sEMG and PSD showed heterogeneous trends across the muscles involved in the task (Fig. 2b). No consistent pattern was observed when comparing muscle activity in the pre-exposure affected, post-exposure affected, and less-affected limbs across the subjects and muscles, as explained below.

The PSD of the less-affected limb trended higher than that of the pre-exposure affected limb for four muscles (FDI, APB, ECRL, EDC) and trended lower than of the pre-exposure affected limb for the other four muscles (FDS, FCU, UT, LT), as seen in Fig. 2b. Also, for some muscles, the PSD decreased from pre-exposure to post-exposure on the affected limb (Fig. 2b, particularly FDI, APB, and ECRL muscles), making it difficult to interpret these trends.

Only the APB, ECRL, and LT muscles demonstrated significant differences across the pre-exposure affected, postexposure affected, and less-affected limbs using PSD and RMS metrics (Fig. 3) (*Friedman* 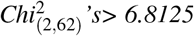, *p’s* < *0.05*). There was a decreasing trend in activation of the LT muscle on both RMS and PSD from pre-exposure affected to post-exposure affected, to less-affected limbs, whereas for the APB and ECRL muscles, the post-exposure affected limb trended to have the smallest RMS and PSD. Conformity scores (i.e., the ratio of the subjects that followed the median group trend to all subjects) indicated that ~30% of the subjects followed the median group trend for RMS and PSD even though statistically significant differences were noted. As we can see in Fig. 4, the ensembled medians of RMS and PSD do not provide any significant trend and the Friedman tests also fail to recognize group effects between the pre-exposure affected, less affected and post-exposure affected limbs (RMS: *Friedman* 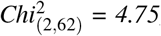, *p* = *0.0930*, PSD: *Friedman* 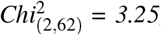, *p* = *0.1969*). Only 22% of the subjects conformed with the the ensembled RMS and PSD trends. Thus, while the classical spectrotemporal analyses of sEMG reveal potential changes in specific muscle activities from pre-to-post exposure to sensorimotor information from the less-affected limb, the trends are not consistent across all muscles, and may not provide information about coordination across muscles involved in the task.

**Figure 3.**
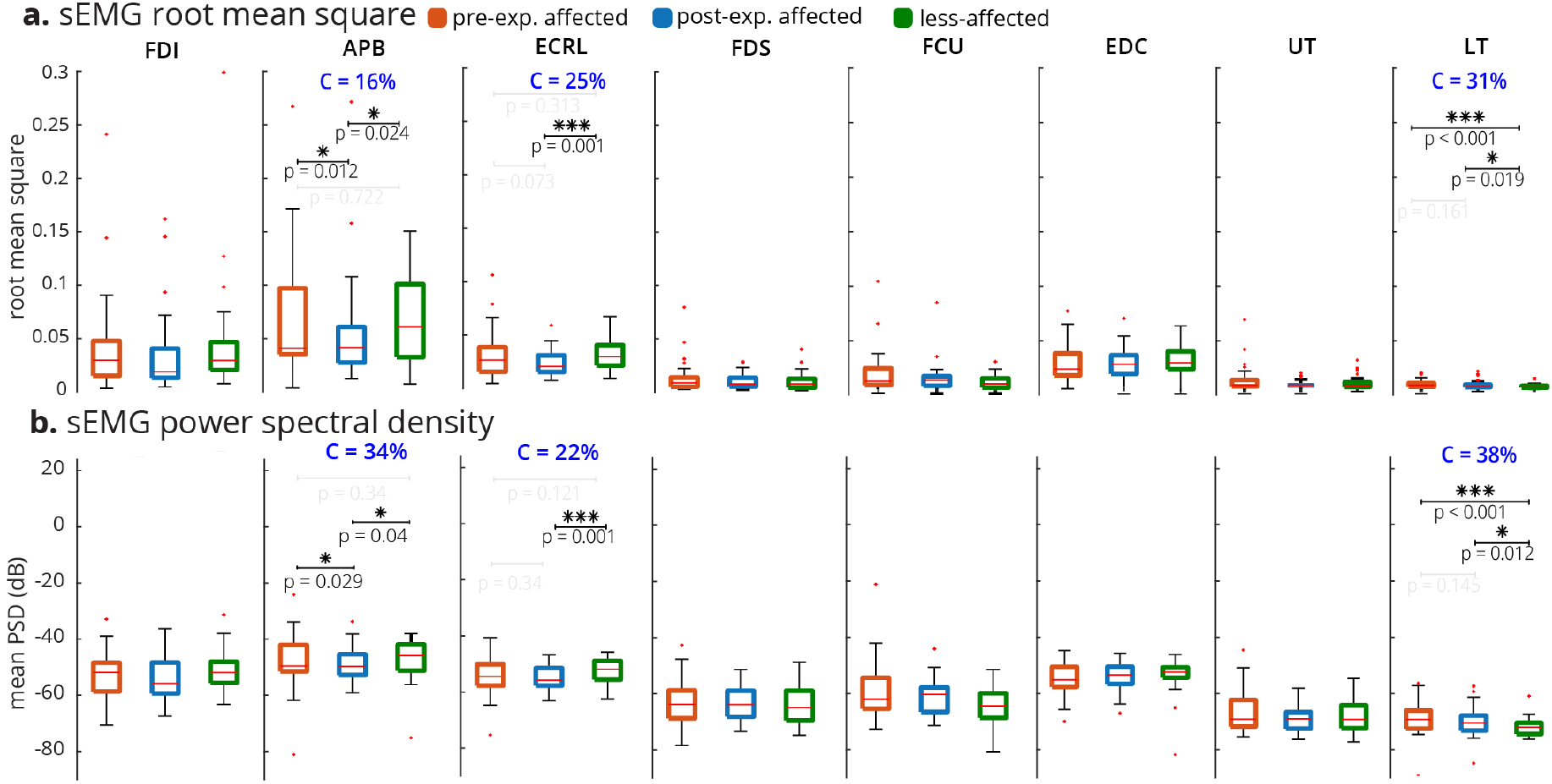
RMS and PSD of sEMG across n=32 subjects. **a.** For each electrode, the median RMS of sEMG envelope is quantified across all trials, and conditions per subject. Differences in the RMS values were only significant for the APB, ECRL, and LT muscles based on the Friedman test. The conformity ratio (the percentage of the subjects that followed the median group trend) for the significant comparisons was C < 31%. **b.** For each electrode, median of average PSD is quantified across all trials and conditions per subject. The differences in PSD were significant for the APB, ECRL, and LT muscles based on the Friedman test. The conformity ratio for the significant comparisons was C < 34%. *** and * indicate significant Friedman and post-hoc Wilcoxon signed-rank test with different p orders.

**Figure 4.**
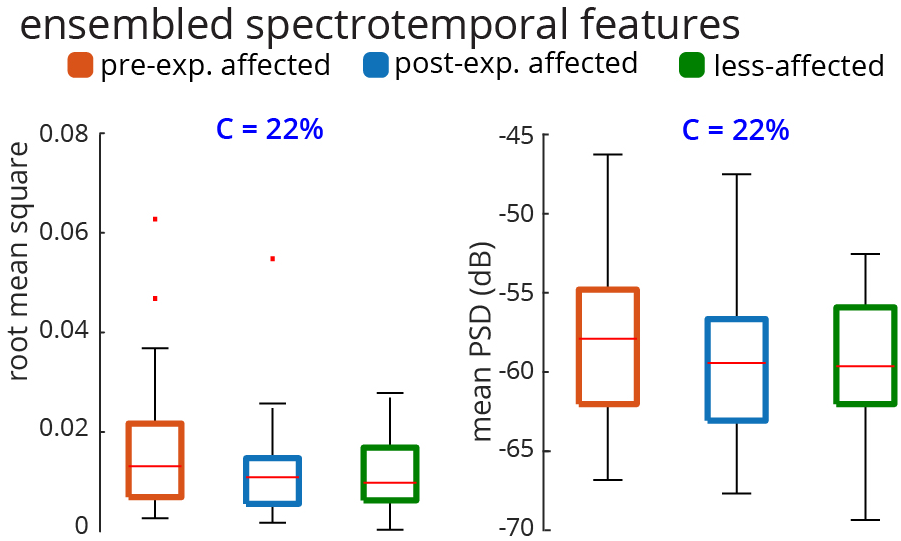
Ensembled RMS and PSD across n = 32 subjects. For each subject, the median RMS values across all sEMG electrodes, all conditions, and all trials are quantified. Thus, there is a distribution of 32 ensembled RMSs for 32 subjects. The same method is used for generating the distribution of ensembled PSD (considering 20 to 200 Hz). The Friedman test was not significant for either RMS or PSD of sEMG (RMS: *Friedman* 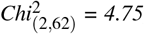, *p* = *0.0930*, PSD: *Friedman* 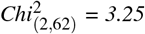, *p* = *0.1969*). The conformity ratio was equal for both RMS and PSD (C = 22%).

### InfoMuNet Analysis

InfoMuNet was used to evaluate the nonlinear coupling of the eight muscles engaged in the functional task. Figs. 1e and 5 show the median connectivity of the functional muscle network across the subjects, where for each subject the network was generated by computing the mutual information exchanged between each muscle pair. As expected, the results from the merged population indicate that the median mutual information degree in the functional muscle network increases monotonically from pre-exposure affected, to post-exposure affected, to the less-affected limbs, highlighting the functional importance and ease of interpretability of this method (Fig. 5a). The mutual information heatmap shows that the post-exposure affected limb demonstrated stronger pairwise coupling than the pre-exposure affected limb, especially for the FCU-FDS, EDC-ECRL and FDS-ECRL muscle pairs involved in grasping and lifting using wrist extension, and in the UT-LT muscle pair involved in stabilizing the shoulder girdle (Fig. 5a). Importantly, the less-affected limb shows even stronger pairwise mutual information coupling than the post-exposure affected limb in these muscle pairs. Overall, the mutual information network became stronger pre-to-post exposure to the sensorimotor information from the less-affected limb (Figs. 1e and 5).

**Figure 5.**
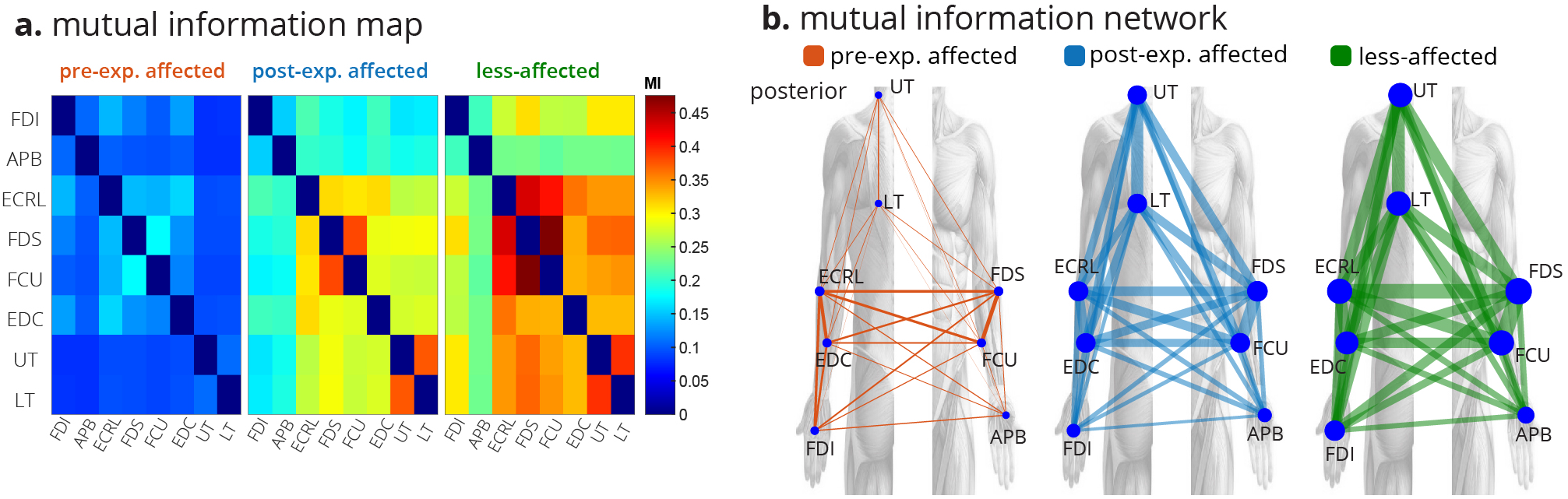
The median InfoMuNet (i.e., the connectivity network of median mutual information between muscle pairs) across all trials of all subjects is shown in two different formats. **a.** The mutual information heatmaps across all subjects show that the pairwise mutual information increases across all nodes (muscles) from the pre-exposure affected to post-exposure affected and less-affected limbs. **b.** The mutual information networks also show an increase in both pairwise mutual information (the edges’ line widths) and the muscles’ degrees (the nodes’ radii).

As can be seen in Fig. 6, the mutual information degree (defined in the Methods section, a measure of mean connectivity of each muscle) of each muscle showed significant increases from the pre-exposure affected to post-exposure affected, to less-affected limbs (*Friedman Chi^2^* > *50.8125*, *p* < *0.001*, *post-hoc p’s* < *0.001*). Overall, high conformity was observed, and interestingly ≥ 94% of the subjects conformed with this trend for the trapezius muscles (UT and LT).

**Figure 6.**
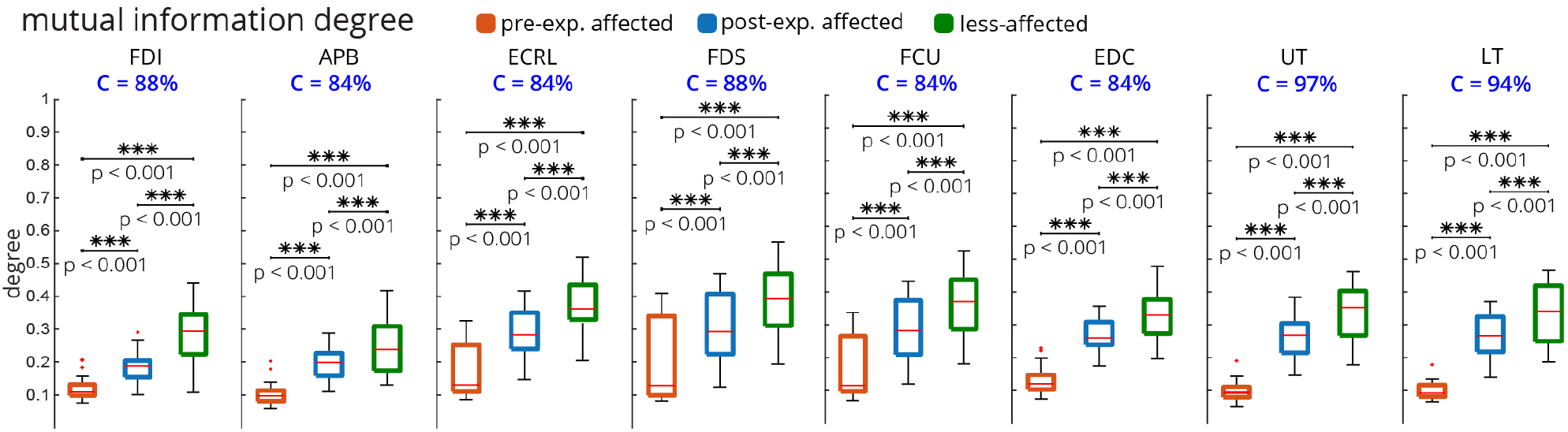
Mutual information degree for individual muscles across all n=32 subjects. For each subject, the mutual information degree was quantified for the median muscle network across all trials. All muscles showed a significant monotonic increase in degree from pre-exposure affected to post-exposure affected and less-affected limbs (*** and ** indicate significant Friedman and post-hoc Wilcoxon signed-rank test). The conformity for all muscles was >80%. The UT and LT muscles had the highest conformity at ≥ 94%.

Furthermore, additional network metrics also showed similar consistent trends from pre-exposure affected to postexposure affected to less-affected limbs (Fig. 7). The mean mutual information degree (averaged across all muscles), overall showed 94% conformity. The mean WCC (defined in the Methods section, a measure of the extent to which the nodes, i.e. muscles, tend to group together), also increased significantly and consistently from pre-exposure affected, to postexposure affected, to less-affected limbs and showed perfect conformity of 100%. The mean shortest path (defined in the Methods section, a measure of network sparsity), decreased from the pre-exposure affected to post-exposure affected to less-affected limbs, with 94% conformity. In addition, the global efficiency (defined in the Methods section, a measure of overall network connectivity), increased from pre-exposure affected to post-exposure affected to less-affected limbs also with perfect conformity of 100%.

**Figure 7.**
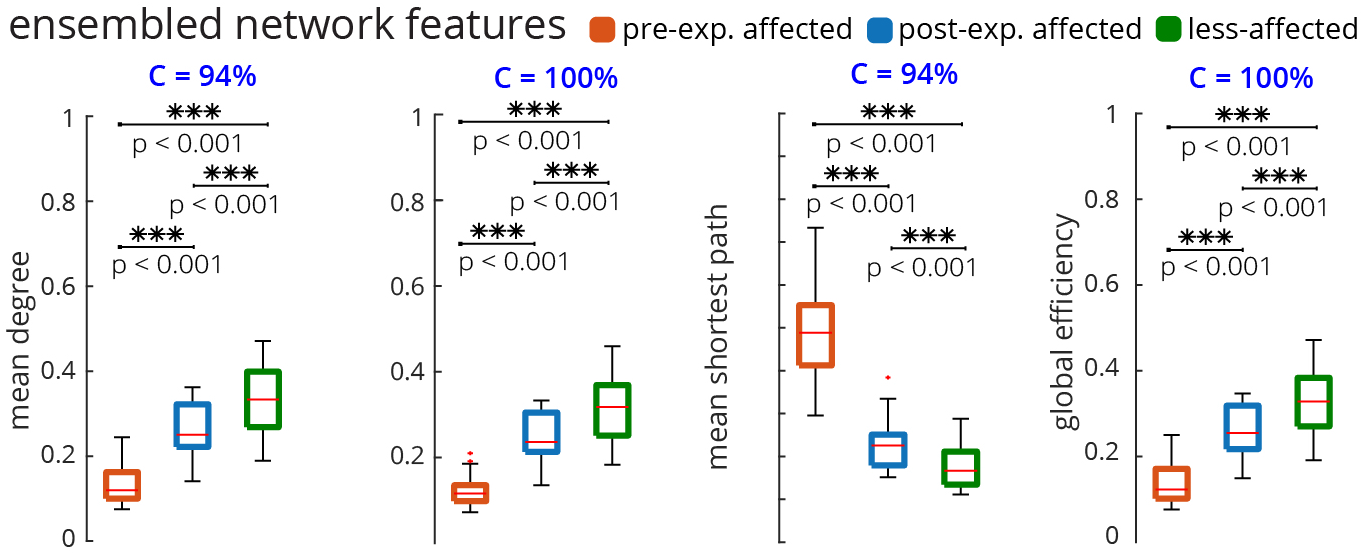
The ensembled metrics of the mutual information network across all n=32 subjects. For each subject, the median muscle network across all trials was computed. Then, the four metrics shown were computed for this ensembled network. All network metrics monotonically changed from pre-exposure affected to post-exposure affected and less-affected limbs (*** indicates significant Friedman and post-hoc Wilcoxon signed-rank test). The mean WCC and global efficiency had perfect conformity, C = 100%. WCC: weighted clustering coefficient

We also investigated the effect of sensory manipulation on the topological changes in the InfoMuNet. Availability of tactile, kinesthetic, and visual sensory inputs changed the mean degree, conformity, and difference between the postexposure affected and less-affected limbs (Fig. 8). The mean mutual information degree increased from the pre-exposure affected, to post-exposure affected, to less-affected limbs for all sensory conditions (*Friedman Chi^2^* > *41.8125*, *p* < *0.001*, *post-hoc p’s* <*0.002*), as seen in Fig. 8a. The conformity was the highest at 97% with the lack of any sensory feedback (all blocked, Fig. 8a), suggesting that subjects had the most consistent behavior under this condition. The conformity consistently decreased with more sensory information was added and was the least at 75% when all tactile, kinesthetic, and visual sensory inputs were provided; this decreasing trend in conformity suggests that inter-individual differences exist in the availability and processing of multiple sensory inputs carried from the less-affected limb, which in turn, influences the functional muscle network connectivity in the affected limb after exposure. The importance of sensory information for muscle coordination was quantified by the ratio of the mutual information degree of the post-exposure affected limb to that of the less-affected limb. A mean ratio of 1 implies that the post-exposure affected limb has similar network behavior to the less-affected limb. In the presence of tactile, kinesthetic, and visual sensory feedback, the ratio was significantly higher (i.e., closer to one) than in the absence of sensory feedback (Fig. 8b) *(Wilcoxon sign-rank test, p = 0.004)*. This result highlights the importance of transfer of sensory input from the less-affected to the affected hand and its role in improving functional muscle network connectivity, which was robustly captured by the InfoMuNet approach.

**Figure 8.**
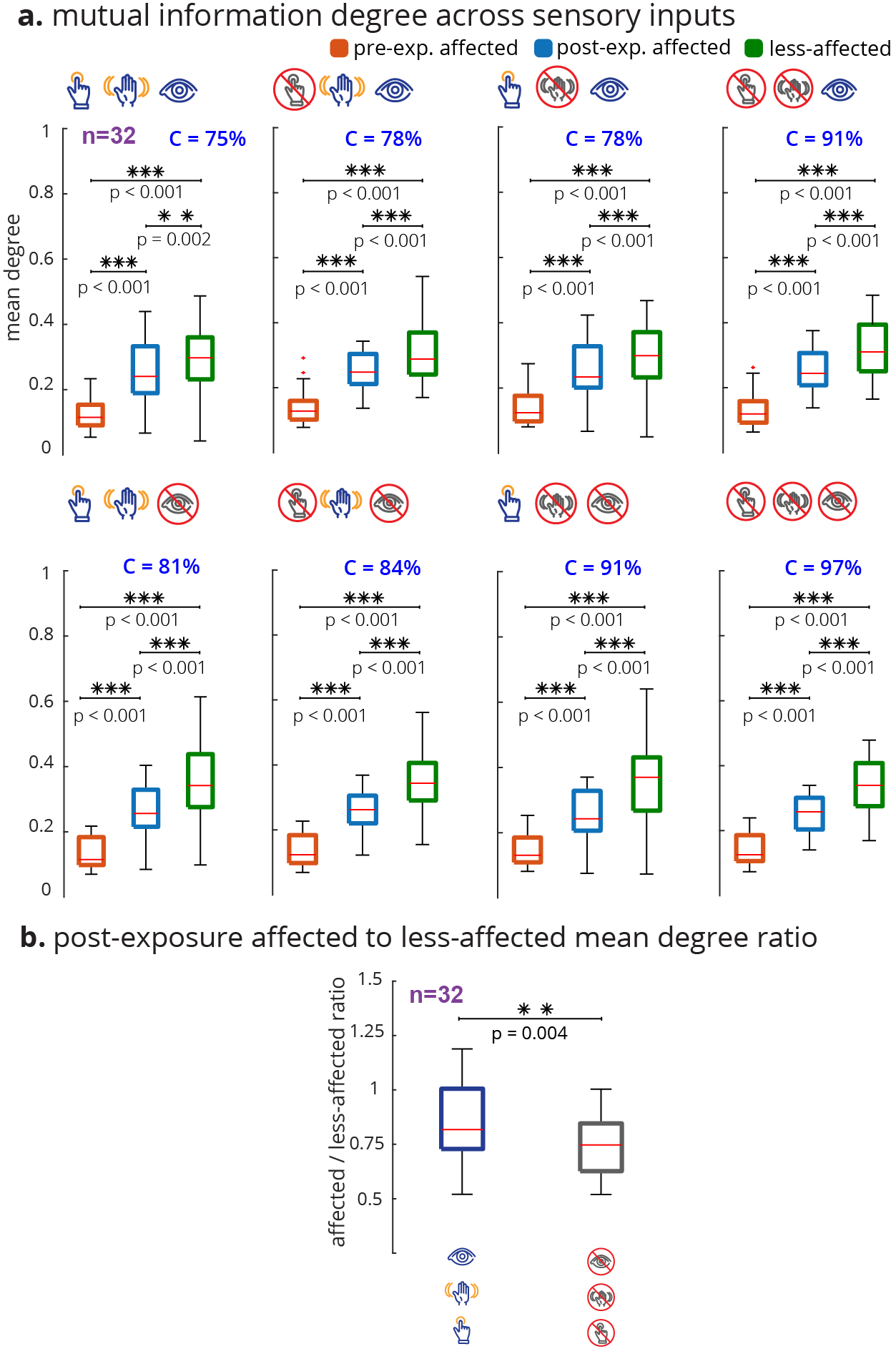
The mutual information degree across sensory conditions. For each subject and sensory condition, the median muscle network across all trials was computed. The mean degree was then calculated for that median muscle network. **a.** For all conditions, the mean degree ascended monotonically with statistical significance from pre-exposure affected to post-exposure affected to less-affected limbs (*** and ** indicate significant Friedman and post-hoc Wilcoxon signed-rank test). **b.** The ratio of the post-exposure affected to less-affected mean degree distributions was computed for all (none blocked) vs no (all blocked) sensory feedback conditions. Wilcoxon sign rank test shows a significant difference between the mean degree ratio distributions. The affected limb’s muscle network approached that of the less-affected limb with sensory feedback compared to without sensory feedback.

## Discussion

The results show that the information theory-based muscle network (InfoMuNet) can encode nonlinear functional muscle coordination as a robust biomarker of sensorimotor integration after stroke with >90% conformity of the individuals to the groups trends. For the first time, we show that the contralateral sensory exposure in chronic stroke patients can result in significant improvement of muscle coordination, quantified by InfoMuNet. This demonstrates the importance and potential for sensory-based rehabilitation and the possibility of improving muscle coordination in the chronic post-stroke population. The functional muscle network also exhibits high sensitivity to the contribution of specific tactile, kinesthetic, and visual sensory inputs for motor coordination. Our results highlight the role of sensory inputs from the less-affected limb to improve functional muscle coordination in the affected limb post-stroke, using the AHT protocol. On the other hand, the spectrotemporal characteristics of sEMG signals fail to capture such changes in muscle activity in a consistent manner. The results highlight the importance of the proposed functional muscle network for tracking progress towards recovery during rehabilitation.

The analysis of sEMG activity showed that muscles have a wide range of temporal and spectral variability during the grasp-and-lift task. The ECRL, EDC, and FDI muscles had the highest sEMG range in the posterior aspect of the limb, and the APB had the highest sEMG range among the anterior muscles (Fig. 2). Conventional temporal (RMS) and spectral (PSD) characterizations of sEMG activations show that specific muscles, specifically the Abductor Pollicis Brevis and Extensor Carpi Radialis Longus muscles demonstrate significant changes in activation from pre-to-post exposure in the affected limb, consistent with their importance in sensing object weight during the grasp-and-lift task as shown previously^32, 33^. However, the conformity of subjects to the ensmebled median group trend was only 22% and the ensembled group trend was not statistically significant, indicating that subjects demonstrated heterogeneous spectrotempporal trends (Fig. 4). The conventional metrics also produced inconsistent relationships across muscles (as seen in Fig. 3), suggesting that spectrotemporal features are not well-suited to use as a biomarker for coordination across muscle groups.

However, as hypothesized, we show for the first time that the InfoMuNet method provides a consistent, uniform gradient (Figs. 5, 6, 7, 8) which highlights its potential as a robust functional biomarker of coordination across muscle groups. The monotonicity of InfoMuNet was validated by computing the mutual information degree of network connectivity, and the results showed a consistent improvement in functional muscle network connectivity from pre-to-post exposure in the affected limb and high discriminative power in distinguishing between the less-affected and post-exposure affected muscle networks. The objective measures of connectivity (e.g., degree and WCC) revealed a consistent trend, where the less-affected limb always had a better score when compared to the post-exposure affected limb. Similarly, the post-exposure affected limb demonstrated a better score compared to the preexposure affected limb across all muscles (Fig. 6); and the trend was consistent for the overall network as well (Fig. 7). Thus, for the first time, this paper shows that the connectivity between the muscles calculated using mutual information (i) is highest for the less-affected limb compared to the affected limb, (ii) increases in the affected limb after sensorimotor exposure to the less-affected limb, and (iii) provides ≥ 94% average conformity across the subjects (Fig. 7). To the best of the authors’ knowledge, this is the first time that information theory has been used to generate a functional muscle network. This is also the first time that the functional muscle network has been used to track changes in sensorimotor integration after stroke.

Stroke tends to increase the variability in the firing rate of the alpha motor neurons^34^, which may explain why nonlinear coupling between muscles is lower in the affected limb. The high conformity of the subjects to the group trends and the high gradient uniformity of the trajectory across the less-affected, post-exposure affected, and pre-exposure affected limbs suggests that InfoMuNet can be used as a biomarker to distinguish differences between the affected and less-affected limbs, and track progress during rehabilitation and recovery. This paper provides the first evidence that, in contrast to the weak and heterogeneous performance of conventional metrics, InfoMuNet is a robust method to track motor coordination during functional task performance as shown here in the context of a sensorimotor exposure paradigm. Moreover, since subjects were not selected based on location of the lesion, it can be concluded that InfoMuNet was successful in capturing the changes in muscle connectivity despite the inherent heterogeneity in the stroke population.

The subjects in this study were in the chronic stage poststroke and were not receiving active rehabilitation at the time of the study. Furthermore, the testing sessions were performed within a few days of each other. Hence, the changes in functional muscle connectivity cannot be attributed to natural recovery. Thus, sensorimotor exposure to the less-affected limb is the likely cause of improved network performance in the affected limb. In a previous study we showed that sensorimotor exposure to the less-affected limb can improve fingertip force coordination in the affected limb after stroke^12^. However, this is the first time that changes in functional muscle network connectivity have been examined. Improved motor coordination in the affected limb after exposure to the less-affected limb may be due to transfer of information at the spinal or supraspinal levels. Previous studies have shown that central pattern generators (CPGs) may be responsible for transfer of learning across limbs^35–37^. CPGs can have spinal or supraspinal origins, hence further research is needed to determine the mechanism underlying this improvement^38^.

In this paper, we also investigated the differential effect of providing tactile, kinesthetic, and visual sensory inputs (Fig. 8). The results show that InfoMuNet is responsive to changes in sensory feedback and reflects how sensory information is used for muscle coordination during sensorimotor integration. It should be highlighted that as the sensory conditions changed, the difference in functional muscle network connectivity captured by InfoMuNet was statistically differentiable across the affected and less-affected limbs (Fig. 8). This further confirms the role that sensory inputs play in motor coordination, and InfoMuNet is robust enough to discriminate the effect of specific sensory inputs on task-specific functional motor coordination after stroke. Examining the ratio of the mean degree of connectivity between the affected and less-affected limbs between all-sensory vs no-sensory feedback showed that the muscle network of the affected limb significantly improves towards that of the less-affected limb with the addition of sensory feedback (Fig. 8b). While the motor task could still be performed in the no sensory feedback condition, sensory feedback augmented sensorimotor control and task-specific intermuscular coordination, further reinforcing the use of InfoMuNet as a biomarker of sensorimotor integration.

Muscle synergy analysis is another approach to quantify the activation patterns across muscles, in which an estimated low-dimensional vector of muscle weights (W) and a corresponding vector of time-varying activation coefficients (A) build the overall muscle activity^39^. The complexity of W can differentiate the motor skills and different control strategies, while A is indicative of how much each element of W is recruited for a given task. Synergy analysis has been successful in differentiating skilled versus non-skilled actions, various postural balance conditions, healthy subjects and stroke patients^40–42^. However, previous studies did not indicate changes in W caused by stroke in the patient group (for example one W for the affected and another W for the less-affected limb). In contrast, previous studies supported the notion of having the same W’s for both affected and less-affected limbs in stroke patients and emphasized the changes only in the activation coefficients (A)^23, 43^. This suggests that synergy reflects mainly spinal control^23^ and does not indicate supraspinal changes associated with stroke. Hence, muscle synergy analysis may not be a robust biomarker for tracking progress in stroke rehabilitation and recovery.

The strong statistics, high conformity, and high gradient uniformity of the results using InfoMuNet suggest that mutual information-based connectivity approach can be used as an objective biomarker for monitoring recovery. Over 90% of the subjects conformed to the median group network trends, suggesting that the InfoMuNet metrics are strong indicators for functional motor coordination after stroke and can sensitively capture changes during the course of recovery (Fig. 7). It should be highlighted that the InfoMuNet metrics had substantially greater conformity than the RMS or PSD metrics and were robust to the inherent intersubject variability present in patients with stroke. Given that existing measures of motor function after stroke do not take sensory feedback into account^14, 15, 44, 45^, InfoMuNet offers quantification of sensorimotor integration, not just motor capability, which is critical for coordinated functional performance. Major advantages in using InfoMuNet for analysis of muscle activity are the objectivity of the metric, ability to test sensorimotor integration and the potential to provide real-time feedback on task performance.

This study examined a heterogeneous group of subjects with chronic stroke at a cross-section in time, precluding an understanding of the effect of time and intervention on poststroke recovery of sensorimotor integration. Furthermore, subjects presented at various times post-stroke. We also did not control for the side of stroke. Previous studies have suggested that hemispheric specialization and lesion location can affect motor control^8^. However, despite this heterogeneity, InfoMuNet emerged as a robust functional biomarker which speaks to its broadest applicability. In future studies we will be able to tease out the effect of stroke-specific characteristics on motor coordination.

In conclusion, InfoMuNet is proposed as a novel, robust functional biomarker of muscle network connectivity required for task performance. It can be used to assess progress in sensorimotor integration and functional performance in patients post-stroke. The results of this paper shed light on the importance of sensory-based motor training to promote healthier coordination of muscle activation. This is a significant observation for chronic post-stroke patients. In a grasp-and-lift task, InfoMuNet successfully differentiated intermuscular coordination in the affected limb after sensorimotor exposure to the less-affected limb. The objective nature of the network metrics and the >90% conformity of individual subjects to the median group trend makes InfoMuNet a strong candidate for future closed-loop assessments of rehabilitation techniques, as it can evaluate both sensory and motor effects on task performance. Its responsiveness to sensory information makes it highly sensitive to even single-session rehabilitation interventions that may predict longer-term responsiveness to interventions, laying the foundation for precision rehabilitation. Future studies on functional muscle network characteristics in individual subjects and their responsiveness to short-term interventions, as well as their long-term effects, will help delineate the underlying neurophysiological processes that influence the recovery of functional muscle networks after stroke and other supraspinal neurodegenerative conditions.

## Methods

Thirty-two patients with chronic stroke (11 females, 21 males, 57.91 ± 12.7 years, mean time since stroke: 46.4 ± 57.5 months, 15 with right hemiparesis and 17 with left hemiparesis) participated in the study after providing informed consent approved by the institutional review board of the New York University (S12-03117). The inclusion criteria were: ability to read/write in English, age >18 yrs, radiologically verified stroke > 4 months old, moderate arm motor impairment (Fugl-Meyer Scale < 60/66), ability to reach, grasp and lift the test objects with the affected side as assessed by the specialist, willingness to complete all clinical assessments and MRI, and comply with training protocols, ability to give informed consent and HIPPA certifications. The exclusion criteria were: sensorimotor impairments in the unaffected hand, severe visual or sensory impairment, including neglect on the affected side, significant cognitive dysfunction (score < 24 on Folstein’s Mini Mental Status Examination), severe or unstable spasticity on treatment with Botulinum toxin or intrathecal baclofen, major disability (modified Rankin Scale > 4), and previous neurological illness, complicated medical condition, or significant injury to either upper extremity.

### Grasp-and-lift task

Subjects completed a series of grasp-and-lift tasks while their muscular activity was recorded (Fig. 2a). The task was to grasp a custom-made instrumented device using the thumb and index fingers and lift it by extending the wrist. The task was performed under eight randomly assigned sensory conditions that allowed or blocked a combination of tactile, kinesthetic, and visual feedback during the task (Fig. 2b). Vision was blocked using a blindfold. Tactile feedback was altered by placing a layer of foam on the subject’s grasping fingertips. Kinesthetic feedback was limited by blocking movement of the wrist using a splint. The arm was placed on an elevated platform and the subject held the object in the air by grasping the device between the thumb and the index finger.

Subjects performed the study over two separate testing sessions: one where they used the affected hand only and a subsequent session during which subjects performed Alternating Hand Training (AHT), where they first grasped and lifted the object with the less-affected hand followed by the affected hand (post-exposure). Subjects performed 7 repeated trials in two sets, each with a light and heavy weight for all sensory conditions. Using the non-parametric Friedman test, the sEMG data did not show significant differences for the spectrotemporal or network metrics. Therefore, we grouped all the weight pairs and performed the analysis on all trials. In total there were eight sensory conditions, and 14 (7 x 2 = 14) trials per sensory condition amounting to 112 trials each with the affected limb pre-exposure, with the less-affected limb, and with the affected limb post-exposure to sensorimotor information from the less-affected limb per subject.

### Data acquisition and spectrotemporal analysis

A wireless sEMG system (DE2.1 Sensors, Delsys Inc., Natick MA) was used to collect sEMG signals from 16 electrode locations, 8 in each limb (Fig. 2c). The recorded muscles were: Abductor Pollicis Brevis (APB), First Dorsal Interosseous (FDI), Flexor Digitorum Superficialis (FDS), Extensor Digitorum Communis (EDC), Flexor Carpi Ulnaris (FCU), Extensor Carpi Radialis Longus (ECRL), Upper Trapezius (UT), and Lower Trapezius (LT). The sEMG signals were pre-amplified and sampled at 2000 Hz, then normalized to the maximum voluntary contraction of each muscle for a given subject. The maximum voluntary contraction was determined by asking the subject to perform maximal isometric contraction of the given muscle for 3s. The maximum activity over the peak 1s was used for normalization.

Data were processed offline using custom functions in MATLAB R2020b (Mathworks Inc., Natick, MA). Signals were band-pass filtered between 20 and 200 Hz, with notch filters at multiples of 60 Hz. The PSD was calculated using the filtered signal by applying Welch’s method with a Hamming window of length 0.85s and 50% overlap. For Figs. 3 and 4, the mean PSD of sEMG in 20-200 Hz was considered for each trial. The Hilbert transform was applied on the filtered and rectified signal to obtain the sEMG envelopes, which were used for RMS computation. For both RMS and PSD computation, the full task duration was considered.

### The InfoMuNet approach

After band filtering between 20 and 200 Hz (with notch filters at multiples of 60 Hz), the sEMG signals were used to quantify the mutual information and construct the connectivity network (InfoMuNet). Mutual information is a metric derived from information theory, which captures nonlinear couplings between two signals *x*(*t*) and *y*(*t*), by computing their individual and joint entropies.

Before computing the entropy, each signal is normalized over the full task duration and then discretized by using the Freedman-Diaconis^46^ method to choose the number of bins, *N_b_*. The discretized version of *x*(*t*) is denoted as *X*, and discretized *y*(*t*) is *Y*. The probability distribution, *P_X_*(*X_i_*), gives the distribution of the discretized signal X and was computed by constructing a histogram from all time samples per trial, with *N_b_* equally spaced bins. The entropy of *X*, *H*(*X*) - i.e. the uncertainty of *x*(*t*) - is given as:

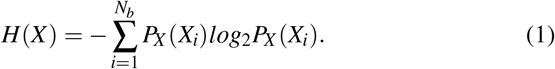

The same method was used to construct the probability distribution for *Y* and hence compute *H*(*Y*), the uncertainty associated with *y*(*t*). The joint entropy, *H*(*X*,*Y*), or joint uncertainty, is inversely proportional to the signals’ inter-dependence and is the final term needed for finding the mutual information. For this, the joint probability distribution for X and Y, *P_X,Y_*(*X_i_*,*Y_j_*) should first be constructed by finding the empirical probability that *X_i_* and *Y_j_* are both in a given bin. Then, the joint entropy, *H*(*X*,*Y*) is given as:

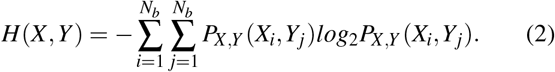

Considering the entropy Venn diagram (Fig. 1d), the mutual information is computed as:

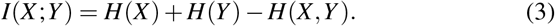

Note that *I*(*X*;*Y*) can alternatively be computed as:

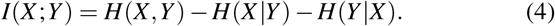

where *H*(*X*|*Y*) and *H*(*Y*|*X*) represent the novel entropies of *X* and *Y* respectively.

Mutual information (*I*(*X*;*Y*)) can be defined as the amount by which a measurement of *y*(*t*) reduces the uncertainty of estimating *x*(*t*)^47^. When *x*(*t*) is completely independent of *y*(*t*), the mutual information is zero. However, if *x*(*t*) and *y*(*t*) are identical time series, the mutual information is maximal.

The median of mutual information for each muscle-pair across trials was quantified, resulting in the median muscle network, which can be represented by adjacency matrix *A*. Using *A*, the following network connectivity metrics were determined: 1) network degree, 2) weighted clustering coefficient (WCC), 3) shortest path, and 4) global efficiency.

1. The (mean) network degree gives an average connectivity for the network. The degree of a node (*D_i_*) is the mean mutual information defining the edges connected to that specific node, as given below:

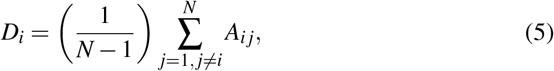

where *N* is the number of nodes. Then, mean network degree, 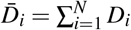, is the mean of all nodes’ degrees.
2. Mean WCC gives the measure of the extent to which nodes in a graph tend to group together. A node’s weighted clustering coefficient (*WCC_i_*) gives a relative measure of how well node *i* is connected to its neighbors (*A_ij_*, *A_ik_*) while also accounting for the neigbors, interconnection (*A_jk_*). A node’s clustering coefficient (*CC_i_*) can be considered the sum of the triangles (∑_*i*_*t_i_*) connected to node i, normalized by the maximum possible value^48^.

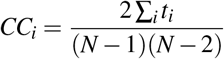 Each triangle’s value will be the product of the three edges, *t_i_* = *A_ij_A_ik_A_jk_*. The weighted adjacency matrix *Ã* is scaled by the maximum connection in the network, hence *Ã_ij_* = *A_ij_*/*max*(*A*). The node’s weighted clustering coefficient *WCC_i_* is then defined as:

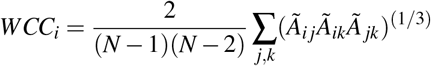 A node which has (i) 0 connectivity to its neighbors or (ii) has neighbors whose interconnections are all 0 will have *WCC_i_* = 0, while a node which is (i) maximally connected to its neighbors and (ii) has neighbors whose interconnections are all maximal has *WCC_i_* = 1. Note that the value of *WCC_i_* is more dependent on node *i*’s connections to its neighbors (*Ã_ij_*, *Ã_ik_* terms) rather than the neighbors’ interconnections (*Ã_jk_* term). The mean WCC, 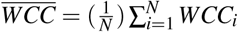, is the mean WCC over all nodes.
3. The mean shortest path, also called characteristic path length, is inversely proportional to how well the network is connected overall. The shortest path between a node pair (muscle *i*, and muscle *j*) is defined as the minimum “connectivity path” (*L_ij_*) between two nodes. A connectivity path between nodes *k* and *l* is 1 /*A_kl_*. However, the shortest path is the minimum sum of paths to travel from *k* to *l*. The shortest path between all nodes in the network was computed using Dijkstra’s algorithm^49^, and the mean shortest path 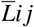 was calculated. The mean shortest path is bounded between 1 (a very well-connected network) and ∞ (a network with ubiquitously zero connectivity).
4. Global efficiency is directly proportional to how well the network is connected overall. The efficiency (*E*) of a network is defined as:

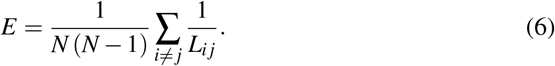 The efficiency (*E*) is normalized by the ideal efficiency (*E_id_*) to give the global efficiency, *GE*:

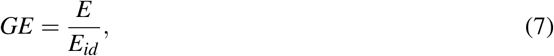

which, unlike mean shortest path, is bounded between 0 and 1. A network with perfect connectivity will have *GE* = 1, while one with no connectivity will have *GE* = 0.

### Statistical analysis

The Kolmogorov-Smirnov test for normality rejected the normal distribution hypothesis for the median PSD, RMS, and connectivity metrics across subjects *(PSD: p*<*0.001 for all of pre-exposure affected*, *post-exposure affected and less-affected*, *RMS: p*<*0.001 for all of pre-exposure affected*, *post-exposure affected and less-affected*, *connectivity metrics: p*<*0.001 for all of pre-exposure affected*, *post-exposure affected and less-affected)*. Therefore, non-parametric statistical tests were used in our analysis. The Friedman test was used to compare spectrotemporal and connectivity measures between the pre-exposure affected, post-exposure affected, and less-affected limbs. The Wilcoxon signed-rank test was used for post-hoc tests if the Friedman revealed significance. Wilcoxon signed-rank test was also used to determine the influence of sensory inputs on the network mean degree ratio. Bonferroni correction was used to account for the multiple comparisons for the post-hoc tests. The significance level for all tests was set at 0.05.

Conformity was quantified as the ratio of the subjects that followed the median group trend. The percentage of the individual subjects that had the same trend as the median group trend was then measured.

To compare the effect of sensory information on the muscle network, the distribution for mean degree ratio of the affected to less-affected sides was computed for the AHT session (Fig. 8b). This is calculated by finding the ratio of the mean degree of connectivity in the post-exposure affected to less-affected limbs, for each of the 32 subjects.

## Disclaimer

This article reflects the views of the authors and should not be construed to represent FDA’s views or policies. The mention of commercial products, their sources, or their use in connection with material reported herein is not to be construed as either an actual or implied endorsement of such products by the Department of Health and Human Services.

